# Bioactive Flavonoid Extract Suppresses NLRP3 Activation and Inflammation in 2D and 3D Lung Models

**DOI:** 10.1101/2025.02.20.639357

**Authors:** Dana El Soufi El Sabbagh, Diulie Valente de Souza, Lauren Pappis, Carolina Bordin Davidson, Maria Mangos, Aizhou Wang, Marcelo Cypel, Liliana Attisano, Alencar Kolinski Machado, Ana Cristina Andreazza

**Author notes:** Corresponding author: Ph.D. Ana Cristina Andreazza. University of Toronto, Department of Pharmacology and Toxicology, 777 Bay St. Toronto, ON, Canada. M5G 2C8.

## Abstract

Inflammasome activation plays a critical role in lung inflammation, with the NLRP3 (NOD-, LRR-and pyrin domain-containing protein 3) inflammasome serving as a key mediator of inflammatory cytokine release and pyroptotic cell death. This study investigates the effects of a bioactive flavonoid extract (BFE), as a potential modulator of NLRP3-mediated inflammation using THP-1 derived macrophages, A549 lung cells (2D cells), and lung organoids (3D cells). Cells were primed with lipopolysaccharide (LPS), treated with BFE, and then activated with nigericin to induce NLRP3 activation and examine BFE’s effects. Markers of inflammation, including presence of ASC specks (3D models), reactive oxygen species production (ROS) (2D models), caspase-1 activity (2D and 3D models), and IL-1β release (2D and 3D models) were measured to assess the extent of inflammasome activation in several treatment conditions. By integrating 2D and 3D lung models, this work provides insight into the NLRP3 inflammasome axis in lung inflammation and explores BFE as a potential therapeutic strategy for inflammasome-driven pulmonary inflammatory processes.

## Introduction

The inflammatory response is a fundamental defense mechanism against pathogens, including bacteria, viruses, and fungi^1^. However, uncontrolled or chronic inflammation can lead to significant cellular damage, particularly in the lungs, where excessive inflammation is implicated in acute and chronic respiratory diseases such as pneumonia, acute respiratory distress syndrome (ARDS), asthma, chronic obstructive pulmonary disease (COPD), and COVID-19^1-3^. A major driver of excessive lung inflammation is the nod-leucine-rich-repeat and pyrin-domain containing protein 3 (NLRP3) inflammasome, a multi-protein complex that plays a pivotal role in innate immune activation and inflammatory cytokine release^4^.

NLRP3 activation occurs in a two-step process: priming (signal 1), where pathogen-associated molecular patterns (PAMPs) or damage-associated molecular patterns (DAMPs) stimulate nuclear factor kappa-light-chain-enhancer of activated B cells (NF-κB) signaling to upregulate NLRP3 expression; and activation (signal 2), where secondary stressors such as reactive oxygen species (ROS), ATP, or nigericin trigger NLRP3 oligomerization, leading to the activation of caspase-1 and subsequent release of IL-1β and IL-18, key mediators of inflammatory responses^5-7^. Dysregulated NLRP3 activity has been implicated in the pathogenesis of inflammatory lung diseases, making it a promising therapeutic target^4,8^.

Current anti-inflammatory treatments, such as corticosteroids, primarily alleviate symptoms but are associated with significant side effects and do not directly modulate NLRP3 inflammasome activity^4^. Small-molecule inhibitors of NLRP3, such as MCC950, have shown therapeutic potential but are limited by toxicity concerns and off-target effects, highlighting the need for safer alternatives^9,10,11,12,13^.

Some phytochemical compounds with anti-inflammatory properties have gained attention as potential modulators of the NLRP3 inflammasome. Our group has been investigating Bioactive Flavonoid Extract (BFE), a flavonoid-enriched fraction derived from *Euterpe oleracea Mart*. (açaí fruit), which contains potent antioxidant and anti-inflammatory molecules, including catechin, epicatechin, and epigallocatechin^14,15,16^. Previous studies have demonstrated that these bioactive flavonoids exhibit strong NLRP3 binding affinity, comparable to synthetic inhibitors, while exhibiting a favorable toxicity profile^17,18^. Furthermore, BFE has been shown to attenuate inflammasome-driven inflammation in macrophages and neuronal cells^19,20,21^.

Given these findings, this study aims to evaluate the anti-inflammatory potential of BFE in lung inflammation models, specifically assessing its ability to modulate NLRP3 activation in both 2D lung epithelial cells (A549) and 3D human induced pluripotent stem-cell (iPSC) derived lung organoids (LOs). LOs provide a physiologically relevant iPSC-derived organotypic model, allowing for a more comprehensive assessment of BFE’s efficacy in lung inflammation^22,23^. By utilizing both 2D and 3D models, we aim to establish BFE as a potential NLRP3 modulator and a promising therapeutic strategy for inflammatory lung diseases (Figure 1).

**Figure 1.**
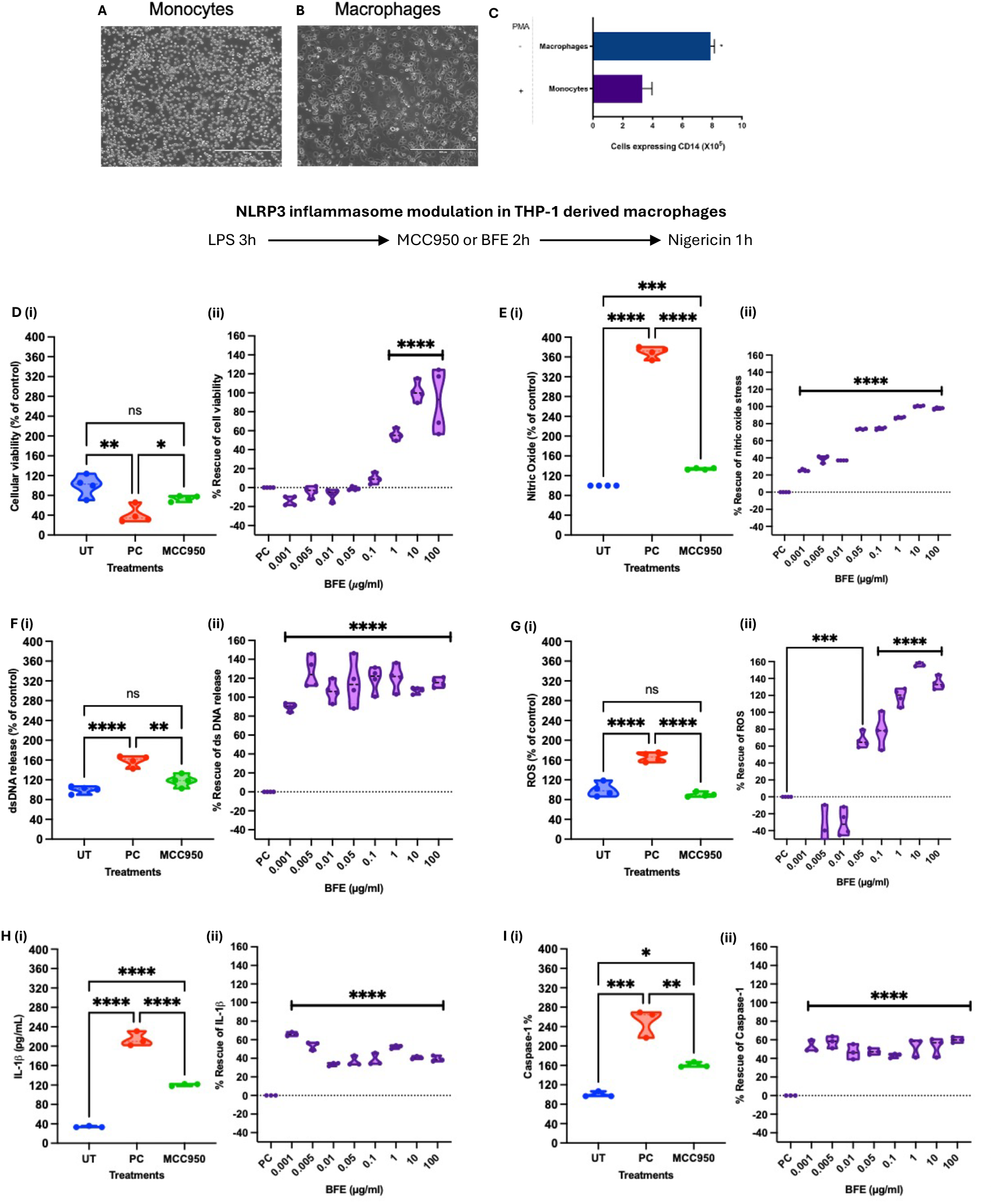
Results of THP1-derived macrophages. A) Representative images of THP-1 monocytes with circular morphology, unattached to the flask. B) Representative image of THP-1 derived macrophages with fusiform morphology, attached to the flask after differentiation. C)Macrophages expressing significantly more CD14 receptors confirming successful differentiation. Scale bar= 50µm. The following conditions were done in figures D-I to (i) validate the system activation using LPS+Nigericin (PC) and MCC950, and (ii) to evaluate BFE’s rescue effects following LPS+Nigericin stimulation, where PC is set as 0% rescue. D) Measurement of cellular viability normalized to untreated (UT) (%); (ii) Rescue effect (%) of cellular viability E) (i) Measurement of nitric oxide (NO) release(%); (ii) Rescue effect (%) of NO following BFE treatment. F) (i) Measurement of dsDNA release; (ii) Rescue effect (%) of dsDNA release after BFE treatment. G) (i) Measurement of reactive oxygen species (ROS) production(%); (ii) Rescue effect (%) of ROS after BFE treatment. H) (i) Measurement of of IL-1β (pg/ml); (ii) rescue effect (%) of IL-1β after BFE treatment. I (i) Measurement of caspase-1 production(%); (ii) Rescue effect (%) of caspase-1 production after BFE treatment. Data expressed as mean ± standard deviation (SD). All statistical analyses were performed using one-way analysis of variance (ANOVA), followed by Tukey’s post hoc test. Values with p<0.05 were considered significant. ** P<0.005 ****P<0.00005

## 2 Materials and Methods

### 2.1 Bioactive Flavonoid Extract (BFE) preparation and characterization

BFE derived from *Euterpe oleracea Mart*. (açaí), was prepared following a previously established protocol by our group^24^. Fresh açaí fruits were obtained from a regulated harvesting area in the Amazon rainforest (3.1019° S, 60.0250° W). The fruits were fully macerated and placed in amber flasks containing 70% ethanol as a solvent at a concentration of 300 mg/mL. The extraction process lasted 21 days, with solvent replacement every seven days to ensure optimal flavonoid yield. After extraction, the material was filtered and concentrated using rotary evaporation to remove residual solvents. The final BFE composition was characterized using high-performance liquid chromatography (HPLC), following the methodology described by Bochi et al ^25^. The detailed chemical matrix determination, including the identification and quantification of key bioactive flavonoids (apigenin, catechin and epicatechin), was previously published by our group ^24^.

### 2.2 2D and 3D cell culture

#### 2.2.1 Use of fibroblasts

A primary human fibroblast cell line was obtained from a healthy volunteer (REB#29949, University of Toronto). Fibroblasts were cultured using Dulbecco’s Modified Eagle Medium (DMEM) (ThermoFischer 11320033) with 10% fetal bovine serum (FBS) (Life Technologies, Burlington, ON, Canada, #10082147) and 1% antibiotics (penicillin/streptomycin) (ThermoFischer 15140122). Cells were kept in a CO^2^ incubator settled to 5% CO2 and 37°C. These cells were used to perform the *in vitro* safety profile tests where cells were exposed to a curve of concentrations of the agent (0.001-1000 µg/mL) for 24, 48 and 72h. After the periods of incubation, cells were evaluated for cellular viability index, nitric oxide (NO) production, ROS levels and double stranded DNA (dsDNA) release described below (supplementary Figure 1).

#### 2.2.2 Generation of THP-1-derived macrophages

THP-1 monocytes (ATCC^®^ TIB-202™) were purchased and cultured with RPMI (Life Technologies, Canada, #11875093) medium, containing 10% FBS and 1% antibiotics (penicillin/streptomycin). Cells were kept in a CO^2^ incubator settled to 5% CO2 and 37°C. THP-1 cells were induced to macrophage differentiation using 25 ng/mL of phorbol 12-myristate 13-acetate (PMA) (ThermoFischer 356150050) in culture for 48 hours ^26^. To confirm macrophage generation, cellular morphology was evaluated from free-floating round cells to a fusiform shape which grow adherent, as well as the CD14 membrane receptor expression by flow cytometry ^27^.

##### 2.2.2.1 Flow cytometry for CD14 membrane receptor expression

THP-1 derived macrophages were collected using TryplE (ThermoFischer 12604013) and resuspended in FACS buffer (1% Bovine Serum Albumin (BSA) in PBS) and counted to a concentration of 10×10^6^ cells in 100/μL. Cells were incubated with Anti-CD14 (Santa Cruz Biotechnology, 5A3B11B5) at 200 μg/ml and an isotype control and incubated for 30 minutes at room temperature. After incubation, cells were washed 3 times and resuspended in a cold FACS buffer then loaded into the Orflow MoxiFLow MXF001 flow cytometer to measure fluorescence.

#### 2.2.3 Culturing 2D Lung cells

A549 cells (ATCC^®^ CCL-185™) were purchased from the Rio de Janeiro Cell Bank. These cells were cultured using DMEM with 10% FBS and 1% antibiotics ww(penicillin/streptomycin). Cells were kept in a CO^2^ incubator settled to 5% CO^2^ and 37°C.

#### 2.2.4 NLRP3 Activation in 2D cells

Both THP-1 derived macrophages and A549 lung cells were plated in 96-well tissue culture plates (Sigma CLS351172) at a concentration of 1×10^5^ cells/mL. After a 24-hour stabilization period, the cells were primed with 100 ng/mL of LPS (Invivogen, San Diego, CA, USA, TLRL-3PELPS LPS) for 3 hours to initiate the NLRP3 inflammasome activation. This was followed by 10μM nigericin (Sigma, St. Louis, MO, USA, N7143), treatment for 1 hour to complete the NLRP3 activation process. To assess treatment efficacy, cells were exposed to either 100nM MCC 950 (Invivogen, San Diego, CA, USA, 210826-40-7) for 2 hours, or a concentration curve of BFE (0.001-100μg/mL) for 2 hours in between the priming and activation steps^28^. Following treatment, the cells were used to measure cell viability and gene expression, while the supernatant was collected and stored at -80°C for subsequent extracellular measurements described below.

#### 2.2.5 Generation of Lung Organoids

LOs were generated at the Applied Organoid Core Facility (ApOC) at the Donnelly Centre, University of Toronto using induced pluripotent stem cells (iPSC) following the protocol published in Mao et al. (2024)^29^. The iPSC line used is a control line purchased from the Gladstone Institutes and is registered in the Human Pluripotent Stem Cell Registry (hPSCReg) under the name UCSFi001-A^30^. In summary, iPSCs were cultured in colonies and then dissociated and plated in V-bottom plates and differentiated into endodermal lineage using DMEM/F12 (Gibco) with supplements including 20% KnockOut Serum Replacement (KSR, Life Technologies), 2% MEM-NEAA, 55 μM β-mercaptoethanol (Life Technologies), 50μM Y-27632, 3 mM CHIR, and 100 ng/mL Activin A. Early stage LOs were then transferred to be grown on air-liquid interface (ALI) using Matrigel (Corning) embedding on membranes to further mature. Media was changed every 3-4 days until they were 6 weeks old and ready for use^29,31^.

#### 2.2.6 NLRP3 Activation in 3D Lung Organoids

NLRP3 inflammasome was activated in the LOs by the methods published in El Soufi El Sabbagh et al. (2024)^32^. Briefly, LOs were incubated with 100ng/mL LPS (Invivogen, San Diego, CA, USA, TLRL-3PELPS LPS) for 3 hours (priming), following 10μM nigericin (Sigma, St. Louis, MO, USA, N7143) for 4 hours (activating). Pretreatment with BFE (1μg/mL) or 100nM MCC950 (Invivogen, San Diego, CA, USA, 210826-40-7) happened in between the priming and activation steps and were 2 hours long. After all incubation periods, LOs were collected and prepared for immunofluorescence of ASC (apoptosis-associated speck-like protein containing a CARD) and supernatant was stores at -80°C for downstream assays.

### 2.2 Experimental assays

#### 2.3.1 Measurement of Cellular Viability

For the 2D cells, cellular viability was assessed using MTT 3-(4,5-Dimethyl-2-thiazolyl -2,5-diphenyl-2H-tetrazolium bromide) (Sigma-Aldrich-M2128; St. Louis, MO, U.S.A.) assay described in Kang et al. (2010). Absorbance was determined at 570 nm using a Synergy H1 (Biotek, Santa Clara, CA, EUA). Since MTT metabolization occurs in viable cells; the decreased absorbance values indicate cell death.

#### 2.3.2 Measurement of Nitric Oxide (NO) levels

Levels of NO were indirectly measured from cell and LO supernatant using a Greiss reagent containing 1% sulfanilamide + 0.1% N-(1-naphthyl)ethylenediamine dihydrochloride. Sulfanilamide reacts with nitrite in the sample to form diazonium salts, which subsequently react with N-(1-naphthyl)ethylenediamine dihydrochloride, producing a purple color^33^. Nitrite levels were quantified using a colorimetric assay based on the Griess reaction. The intensity of the color is directly proportional to nitrite concentration and, consequently, an indirect measure of nitric oxide (NO) production. Absorbance was measured at 540 nm using an Anthos 2010 plate reader (Biochrom®, Anthos 2010, London, UK).

#### 2.3.4 Analysis of double stranded (ds)DNA release

Extracellular dsDNA from the 2D and 3D cell supernatant was quantified using the Quant-iTTM PicoGreen® kit (ThermoFisher, Waltham, MA, USA, P11495) following the protocol outlined by Ahn, Costa, and Emanuel (1996)^34^. PicoGreen binds to dsDNA by intercalating between its strands, emitting fluorescence for measurement. Fluorescence was recorded at 480 nm excitation and 520 nm emission using a plate reader equipped with Gen5 software (BioTek Instruments, Inc., Winooski, VT, USA, 253147), enabling the evaluation of cell integrity.

#### 2.3.5 Semi-quantification of total levels of reactive oxygen species (ROS)

Total levels of ROS were determined in from 2D and 3D cell supernatant using a semi-quantitative method using 2′ 7′ dichlorodihydrofluorescein diacetate (DCFH-DA) (Sigma-Aldrich-D6883; São Paulo, SP, Brazil). DCFH-DA is metabolized by intracellular enzymes to form dichlorodihydrofluorescein (DCFH)^35^. ROS molecules reduce DCFH to dichlorofluorescein (DCF), which fluoresces at 525 nm when excited at 488 nm. Fluorescence intensity was measured using a Spectra iMax i3 plate reader (Molecular Devices, San Jose, CA, U.S.A.)

#### 2.3.6 Gene expression of inflammatory cytokines

The gene expression of IL-1β, IL-6, TNF-α, IL-10, caspase-1, and NLRP3 were measured in A549 cells following the descriptions of the manufacturers. RNA was extracted by using TRI Reagent® (Sigma-Aldrich, Saint Louis, MO, USA) and was quantified using NanoDrop Lite® (ThermoFisher Scientific, Wilmington, DE, USA). The RNA amount was normalized for each sample at a final concentration of 100 ng. The complementary DNA (cDNA) was produced by using the iScript™ cDNA Synthesis kit (Bio-Rad, Hercules, CA, USA) following the manufacturer’s instructions. Quantitative real-time PCR was carried out using a GoTaq® qPCR Master kit (Promega, Madinson, WI, USA). qRT-PCR cycles were as follows: i) 50 °C for 120 s; ii) 95 °C for 120 s; and iii) 40 cycles of 95 °C for 15 s followed by 60 °C for 30 min. IL-1β primers were as follows: forward—5′ AAGCCCTTGCTGTAGTGGTG 3′; reverse—5′ GAAGCTGATGGCCCTAAACA 3′. IL-6 primers were as follows: forward—5′ AGACAGCCACTCACCTCTTCAG 3′; reverse—5′ TTCTGCCAGTGCCTCTTTGCTG 3′. TNF-α primers were as follows: forward—5′ CTCTTCTGCCTGCTGCACTTTG 3′; reverse—5′ ATGGGCTACAGGCTTGTCACTC 3′. IL-10 primers were as follows: forward—5′ GTGATGCCCCAAGCTGAGA 3′; reverse—5′ TGCTCTTGTTTTCACAGGGAAGA 3′. NLRP3 primers were as follows: forward—5′ CCCCGTGAGTCCCATTA 3′; reverse—5′ GACGCCCAGTCCAACAT 3′. Caspase-1 primers were as follows: forward—5′ CGCACACGTCTTGCTCTCAT 3′; reverse—5′ TACGCTGTACCCCAGATTTTGTAG 3′. Beta-actin was the housekeeping gene. Beta-actin primers were as follows: forward—5′ CTGGCACCACACCTTCTAC 3′; reverse: 5′-GGGCACAGTGTGGGTGAC 3′.

#### 2.3.7 Measurement of Caspase-1 expression

Caspase-1 expression was measured using an ELISA procedure previously published by our group^36^. In brief, Supernatant (50 µL) of THP-1-derived macrophages and LOs was plated in a high binding protein plate (Greiner, Monroe, NC, USA, #655061) overnight at 4 °C in a shaker. The next day, wells were washed with PBS-T and blocked with 2% bovine-serum-albumin (BSA) (BSA; Sigma-Aldrich, St. Louis, MO, USA, #A9418) for 1 hour at room temperature. Anti-caspase-1 antibody was used (1:1000 Abcam, Waltham, MA, USA, #ab62698) for 1 hour and secondary anti-body anti-rabbit-HRP (1/1000; Cell Signaling Technology, Danvers, MA, USA, #7074) was incubated for 1 hour at room temperature with washes in between each step. Reaction was developed using TMB solution (Cell Signaling Technology, Danvers, MA, USA, #7004) and stopped with 1M HCl. Absorbance was read at 540nm using a plate reader with Gen5 software (BioTek Instruments, Inc., Winooski, VT, USA, 253147.

#### 2.3.8 Measurement of interleukin-1β (IL-1β)

Human IL-1β production was measured from the supernatant THP-1-derived macrophages and LOs using an ELISA kit from Abcam (2ab214025, Cambridge, UK). Manufacturers protocol was followed, and samples were plated neat. Results are shown in pg/ml. Absorbance was read using a plate reader with Gen5 software (BioTek Instruments, Inc., Winooski, VT, USA, 253147.

#### 2.3.8 Measurement of circulating cell-free mitochondrial DNA (ccf-mtdna)

Levels of circulating cell-free mitochondrial DNA (ccf-mtdna) were measured in LOs following a protocol described in Lu et al (2024)^37^ El Soufi El Sabbagh et al (2024)^32^. Supernatant of the LO was collected (100µL) and mitochondrial DNA was extracted by the QiAMP DNA mini kit. For gene expression, the ND1 and ND4 mitochondrial genes were used, and B2M and PPIA for nuclear genes. All PCR products used are listed in El Soufi El Sabbagh et al (2024)^32^

#### 2.3.9 mmunofluorescence & antibody stains of LOs

LOs were washed with PBS, then placed in 4% paraformaldehyde (PFA) overnight for fixation. The following day they were washed twice with PBS-T and embedded in 30% sucrose overnight in the fridge. They were then washed and embedded in cryomolds in OCT gel and flash frozen with dry ice. LOs were then cryosectioned into 20um slices and kept frozen at -80°C until immunostaining. Primary antibodies used were SPC (1:200) (Invitrogen PAS-71680) and SOX9 (Abcam, ab76997) and ASC (AL177). Secondary antibodies used included Goat anti-Mouse [IgG] [H + L] (Abcam, #Ab97035) secondary antibody Cy3 and Donkey anti-Rabbit IgG [H + L] secondary antibody Alexa Fluor Plus 647, (Invitrogen, #A32795). All primary antibodies were diluted and secondary antibodies (1:750) using 0.5% BSA and after staining, the coverslips were mounted on a glass slide using ProLong Gold Antifade Mountant with DAPI (invitrogen, Waltham, MA, US, P36935). Cryosectioned samples were imaged at the Microscopy Imaging Laboratory, University of Toronto, Canada using an Elyra Super-resolution confocal microscope LSM880.

#### 2.3.10 Measurement of ASC specks in LOs

DAPI positive cells were counted using HALO Indica Labs and HALO AI (v4.0.5107.357) using the analysis tool HighPlex FL (v4.2.3) which counts and characterizes cells in an automated manner. Measurement of ASC specks was done in a semi-automated approach through FIJI (ImageJ) (version 2.9.0/1.53t) and is described in detail in the methods protocol our group published in El Soufi El Sabbagh et al. (2024)^32^.

### 2.4 Statistical Analysis

Results for were plotted in Microsoft Excel (Version 16.77.1) and plotted in GraphPad Prism (version 10.4.1 (532)) for statistical analysis. To assess the effect of BFE, % rescue was calculated relative to the untreated (UT) baseline and to the stimulated LPS+Nigericin positive control (PC) conditions. PC represents the maximal inflammatory response or cellular damage, while UT serves as the healthy baseline reference condition. The % rescue for each BFE dose was determined using the following equation: ((BFE-PC) / (UT-PC) * 100) where BFE’s Value corresponds to the measured response (e.g., cell viability) at a given drug concentration. This calculation expresses the degree to which BFE restored the system toward UT baseline levels. A value of 100% rescue indicates full restoration to UT levels, while 0% rescue indicates no improvement compared to PC. Negative values represent further deviation from the UT condition. Data was analyzed using a student’s t-test or a One-way Anova followed by Tukey post hoc. Comparisons with P<0.05 were considered significant.

## 3. Results

### 3.1 BFE modulates the inflammatory response in THP-1 derived Macrophages

Successful generation of macrophages was confirmed via morphological microscopic analysis demonstrating free-floating round cells to a fusiform shape which grow adherent as shown in Figure 1A and B. Moreover, analysis of CD14 membrane receptor expression by flow cytometry (Figure 1C), demonstrated that the generated macrophages express higher levels of CD14 than monocytes, consistent with prior literature ^38,39^.

#### 3.1.1 Validation of System Activation and Inhibition in THP-1 derived Macrophages

Successful NLRP3 activation was confirmed in THP-1 derived macrophages by measuring direct downstream markers of inflammation like caspase-1 and IL-1*β* levels under three conditions: baseline control (UT), LPS+nigericin positive control (PC), and MCC950 (classical NLRP3 inhibitor) (Figure 1 Hi-Ii, respectively).

For overall cellular health assessments, following PC and MCC950 treatments, cellular viability, NO release, ROS generation, and dsDNA levels were measured. In the PC group, cell viability was significantly decreased, while significantly increasing levels of NO release, dsDNA release, and ROS formation (Figure 1Di-Gi) confirming inflammasome activation-induced cellular stress. PC treatment group significantly decreased cellular viability, and heightened levels of inflammatory markers and cellular stress, indicating successful activation of the inflammasome and treatment with MCC950 partially restored these effects.

Since caspase-1 and IL-1*β* are direct downstream effects of NLRP3 activation, their levels were quantified to confirm inflammasome activation. A significant increase in both pro-inflammatory cytokine levels in the PC group indicated successful inflammasome activation and a decrease of caspase-1 and IL-1*β* levels as shown by MCC950 group, confirmed effective inflammasome inhibition using a known inhibitor (figures 1Hi-Ii). Collectively, we demonstrated the ability to reproduce the inflammasome pathway in THP-1 derived cell model.

#### 3.1.2 Evaluation of BFE’s Rescue Effects in THP-1 derived Macrophages

We next evaluated the therapeutic potential of BFE as a novel inhibitor to NLRP3-mediated inflammation, its % rescue effects on cells were assessed in all the assays outlined above. Treatment of macrophages with BFE at varying doses revealed a significant increase in cellular viability and decrease in ROS formation in a dose dependent manner, particularly at 1-100μg/mL (Figure 1Dii and Gii). Concordantly, increases in NO production, dsDNA release, and il-1*β* and caspase-1 the levels of were also attenuated at all doses of BFE ranging from 0.001-100μg/mL (Figure 1 Ei-Fi, Hi-Ii).

### 3.2 BFE modulates the inflammatory response in A549 Lung Cells

We next examined the effect of BFE on NLRP3 inflammasome using a human lung carcinoma epithelial cell line, A549. A549 lung cells were plated and treated with the three main conditions (UT, PC, MCC950) as well as varying doses of BFE (Figure 2A-D).

**Figure 2.**
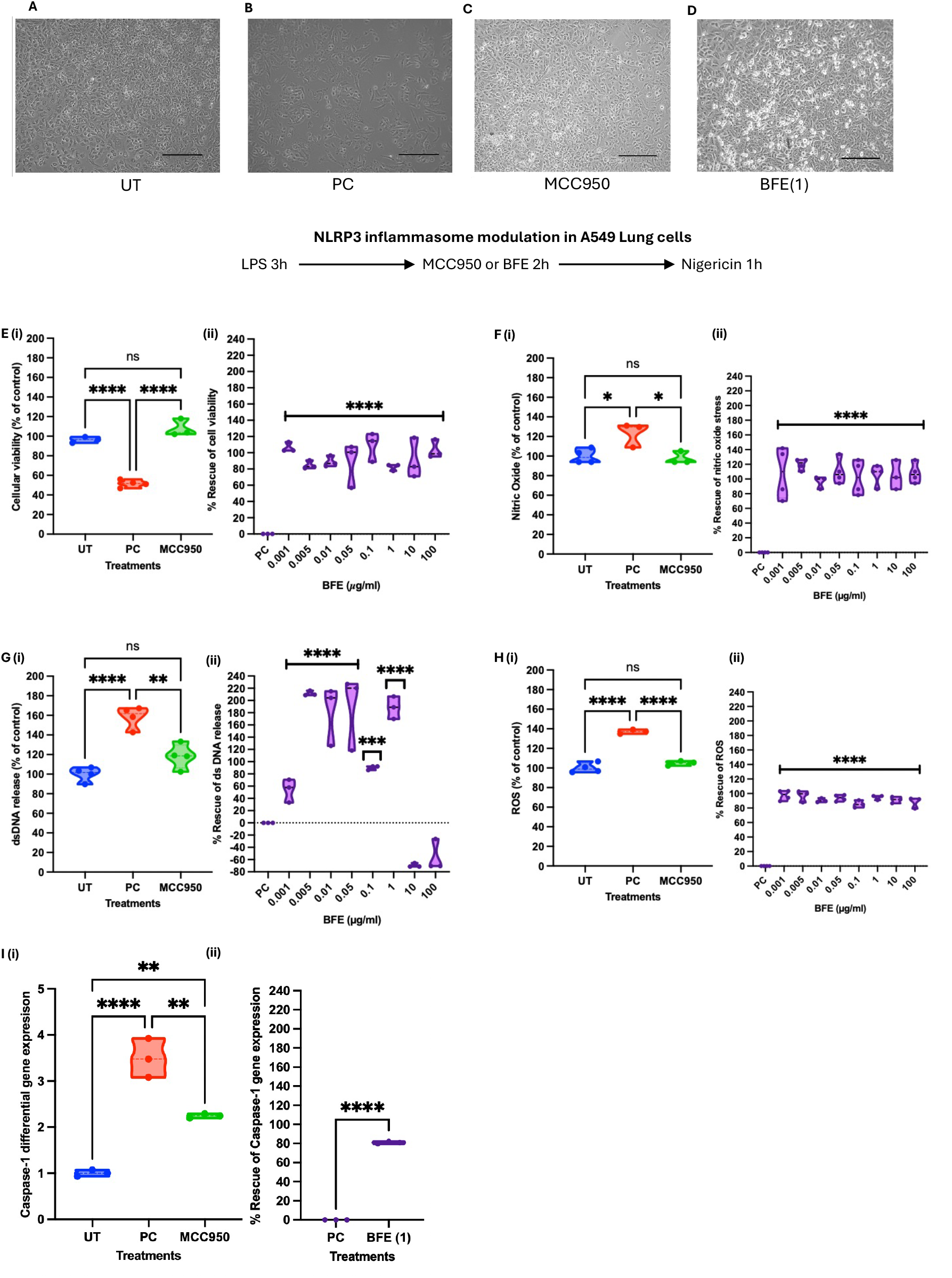
Results using A549 2D Lung cells. Representative images of A)Untreated (UT)): cells in culture medium. B) Positive control (LPS+Nigericin): cells with 100ng/mL of LPS followed by 10nM of nigericin for 1 hour. C) MCC950 control: cells exposed to 100ng/mL of LPS followed by 100nM of MCC950 for 2 hours and then 10nM nigericin 1 hour D) cells exposed to 100ng/mL of LPS followed by BFE (1µg/mL) for 2 hours followed by 10nM of nigericin for 1 hour. Scale bar=50µm. The following conditions were done in figures E-I to (i) validate the system activation using LPS+Nigericin (PC) and MCC950, and (ii) to evaluate BFE’s rescue effects following LPS+Nigericin stimulation, where PC is set as 0% rescue. E) Measurement of cellular viability normalized to untreated (UT) (%); (ii) Rescue effect (%) of cellular viability F) (i) Measurement of nitric oxide (NO) release(%); (ii) Rescue effect (%) of NO following BFE treatment. G) (i) Measurement of dsDNA release; (ii) Rescue effect (%) of dsDNA release after BFE treatment. H) (i) Measurement of reactive oxygen species (ROS) production(%); (ii) Rescue effect (%) of ROS after BFE treatment. I (i) Measurement of caspase-1 gene expression; (ii) Rescue effect (%) of caspase-1 gene expression after BFE treatment. Data expressed as mean ± standard deviation (SD). All statistical analyses were performed using one-way analysis of variance (ANOVA), followed by Tukey’s post hoc test. Values with p<0.05 were considered significant. ** P<0.005 ****P<0.00005

#### 3.2.1 Validation of System Activation and Inhibition in A549 Lung Cells

Cellular viability, NO release, dsDNA production, and ROS formation were assessed under the three conditions: UT, PC, and MCC950 (Figure 2Ei-Hi, respectively). To validate successful NLRP3 inflammasome activation, caspase-1 levels were measured and found to be significantly increased (Figure 2Ii). PC treatment group significantly decreased cellular viability, and heightened levels of inflammatory markers and cellular stress, indicating successful activation of the inflammasome. Treatment with MCC950 significantly restored these effects.

#### 3.2.2 Evaluation of BFE’s Rescue Effects in A549 Lung Cells

To assess the therapeutic effects of BFE, the rescue percentage was assessed its % rescue effects on cells in all the assays outlined above. BFE’s treatment significantly increased cellular viability in a dose-dependent manner (Figure 2Eii). Similarly, NO production, dsDNA release, and levels of ROS formation were rescued using a range of BFE doses, with the most effective dose being 1μg/mL (Figure 2Fii-Hii). Gene expression levels of caspase-1 were also rescued effectively by 85% in the 1μg/mL BFE dose (Figure 2Iii). Additionally, 1 µg/ml of BFE was previously described as an optimal concentration in other studies developed by our group^19-21^.

### 3.3 BFE *shows efficacy in reversing NLRP3 activation in 3D Lung Organoids (LO)*

Our group has previously published on activating the NLRP3 inflammasome *in vitro* using both 2D cells including peripheral blood mononuclear cells, microglia and neuroblastoma cells, and 3D cultures, cerebral organoids^28,32^. Since the focus of this study is on the lung, we next utilized LOs as a sophisticated model which offers a more complex system of multiple lung cell types interacting together. Lung organoids were generated from WTC11 iPSC cells as previously described. The presence of diverse cell types including SOX9 progenitors which play a crucial role in epithelial cell generation and SPC to identify AT2 cells, essential for surfactant protein and alveolar homeostasis in 6-week-old LOs was confirmed (Figure 3A-B).

**Figure 3.**
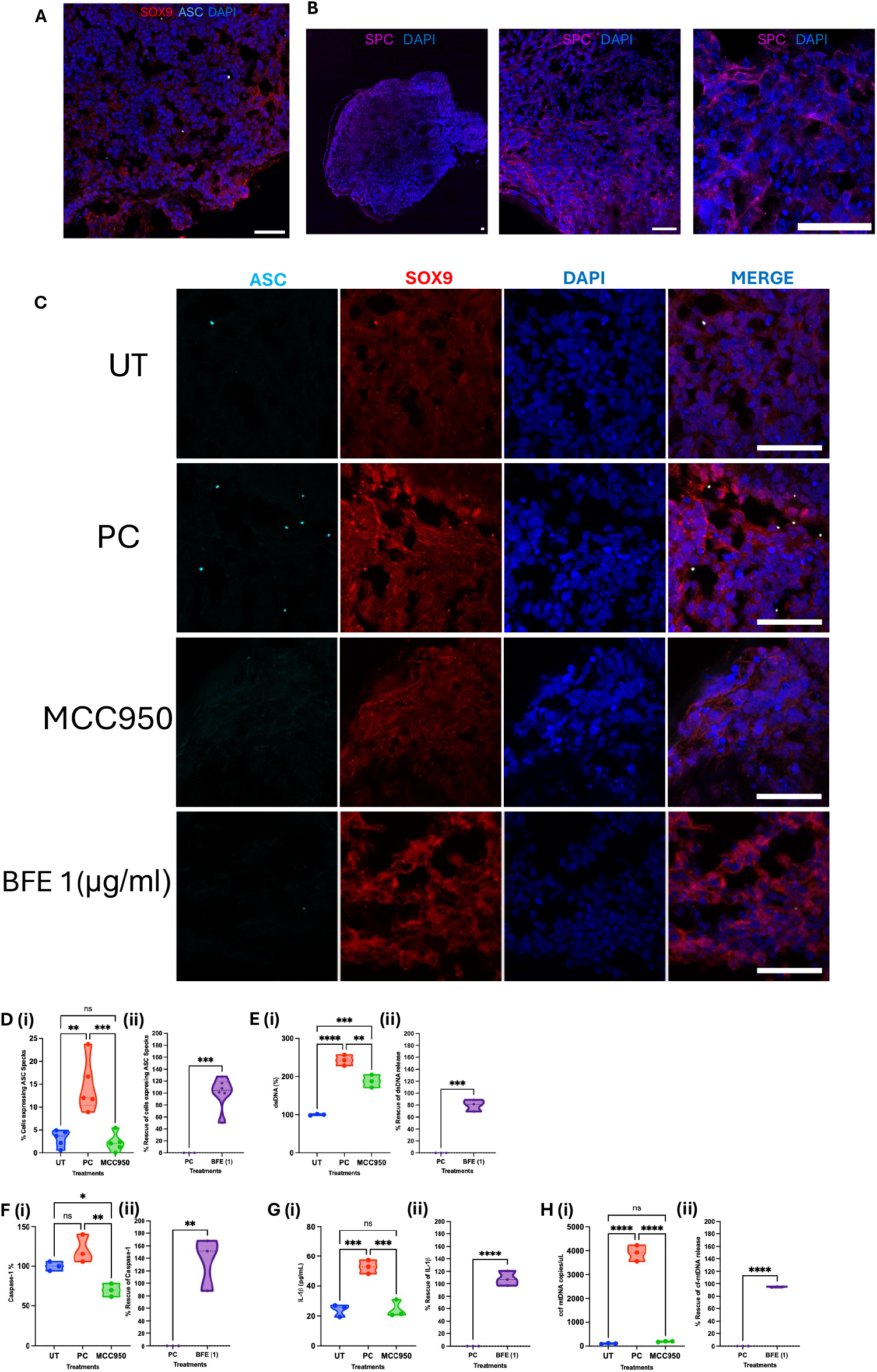
Testing the BFE extract using Lung Organoids. Representative immunofluorescence images of lung organoids at 6 weeks using A) SOX9 positive cells and ASC specks B) SPC protein C) NLRP3 inflammasome activation in 4 treatment groups staining for SOX9, ASC, and DAPI. The following conditions were done in figures D-H to (i) validate the system activation using LPS+Nigericin (PC) and MCC950, and (ii) to evaluate BFE’s rescue effects following LPS+Nigericin stimulation, where PC is set as 0% rescue. D) Measurement of cells expressing ASC Specks (%); (ii) Rescue effect (%) of cells expressing ASC specks after BFE treatment. E) (i) Measurement of dsDNA release (%); (ii) Rescue effect (%) of dsDNA following BFE treatment. F) (i) Measurement of caspase-1 protein expression (%); (ii) Rescue effect (%) of capsase-1 after BFE treatment. G) (i) Measurement of IL-1β (pg/ml); (ii) Rescue effect (%) of IL-1β after BFE treatment. H) (i) Measurement of cf-mtDNA release (%) (ii); Rescue effect (%) of cf-mtDNA after BFE treatment. Data are presented as mean ± standard deviation. Statistical significance was determined using one-way ANOVA with Tukey’s post-hoc test. **represents P<0.005; ***P<0.0005; ****P<0.00005; Scale bar = 50µm.

#### 3.3.1 Visualization of NLRP3 inflammasome Activation and Rescue in LOs

Since this study involves 3D tissue, we applied the same methodology previously developed to activate the NLRP3 inflammasome to examine the effects of BFE in LOs^32^. Inflammasome activation was visualized by detecting ASC (apoptosis-associated speck-like protein containing a CARD) speck formation, which occurs when ASC oligomerizes into discrete cytoplasmic puncta upon NLRP3 activation. These ASC specks serve as a hallmark of inflammasome assembly and were quantified in the PC group, where an increased number of specks confirmed successful inflammasome activation (Figure 3C and 3Di). After confirming NLRP3 activation through elevated number of ASC specks in the PC group we found the BFE to modulate the inflammasome’s effects through decreased number of specks in comparison to the PC (Figure 3Dii).

#### 3.3.2 Evaluation of NLRP3 Inflammasome Activation and Rescue Effects

To further examine BFE’s effects on cellular stress and inflammasome activation, levels of dsDNA, caspase-1, and IL-1β were measured (Figure 3E-G)^5^. Levels of ccf-mtdna were also measured as a marker of overall cellular stress as activation of the NLRP3 inflammasome can lead to pyroptotosis which forces cells to release not only nuclear but mitochondrial DNA into the extracellular space (Figure 3H)^5^. In all four assays, successful activation of NLRP3 inflammasome was shown through a significant increase in levels compared to the UT (Figure 3Ei-Hi).

The treatment of 1µg/ml BFE also showed significant rescue effects to levels of dsDNA, caspase-1, IL-1β, and ccf-mtdna, restoring the cellular homeostasis to levels close to UT (Figure 3Eii-Hii). These results highlight the potential of BFE not only to inhibit NLRP3-induced inflammation and injury but also to provide enhanced cellular protection, offering an advantage over MCC950.

## 4 Discussion

This study demonstrates that our Bioactive Flavonoid Extract (BFE) is capable of suppressing NLRP3 inflammasome activation and consequent inflammatory cytokines release in lung models, using both 2D A549 lung epithelial cells and iPSC-derived 3D lung organoids. Our results show that BFE can rescue the cellular viability, modulates inflammatory responses at multiple levels, significantly reducing ROS and NO production, dsDNA formation, and pro-inflammatory cytokines (IL-1β and caspase-1) release compared to untreated inflammatory controls. Notably, BFE exhibited similar or slightly superior efficacy to MCC950, a well-known synthetic NLRP3 inhibitor, while maintaining a favorable safety profile. These findings suggest that BFE could serve as a natural, low-toxicity alternative for targeting inflammasome-driven lung inflammation.

NLRP3 inflammasome activation follows a two-step process (Figure 4): priming (signal 1) and activation (signal 2)^40,41^. The priming phase is necessary for initiating inflammasome activity by upregulating key inflammasome components, including NLRP3, pro-IL-1β, and pro-IL-18, through NF-κB activation. LPS, a pathogen-associated molecular pattern (PAMP), binds to Toll-like receptor 4 (TLR4) on immune and lung epithelial cells, initiating a signaling cascade that activates MyD88, a critical adaptor protein in the TLR4 pathway^5,40^. This interaction triggers the downstream activation of NF-κB, a transcription factor that is translocated to the nucleus and enhances the expression of NLRP3, pro-IL-1β, and pro-IL-18, along with other inflammasome-related proteins^5,40^. Although priming is essential for inflammasome readiness, it does not induce full activation. Instead, an additional stimulus (signal 2) is required to trigger inflammasome assembly and cytokine maturation.

**Figure 4.**
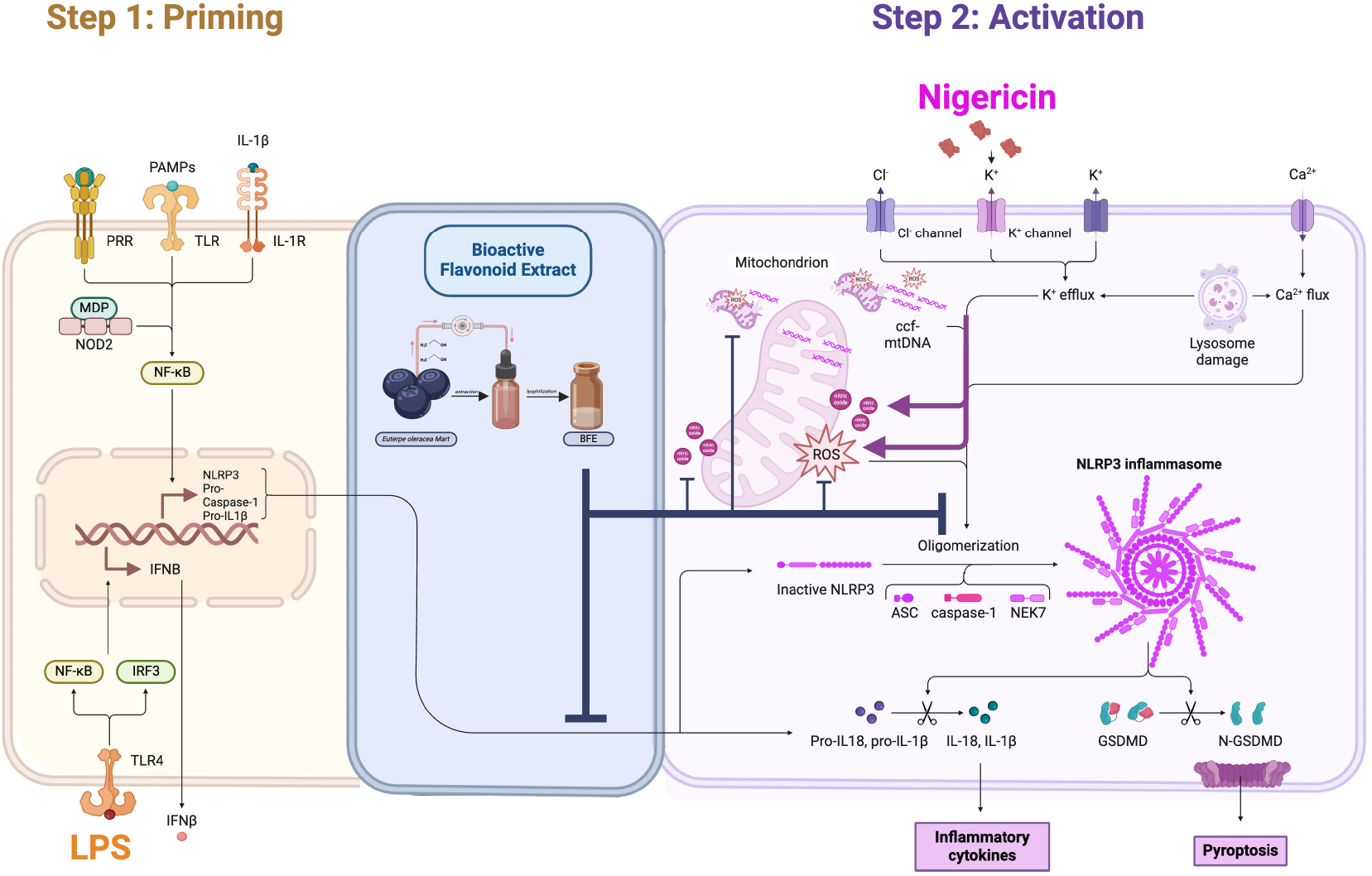
NLRP3 inflammasome activation through LPS and Nigericin treatment and effects with bioactive flavonoid extract (BFE). Step 1 (Priming): Lipopolysaccharide (LPS) binds Toll-like receptors (TLRs) and triggers downstream NF-κB signaling, leading to the transcriptional upregulation of NLRP3, pro-caspase-1, and pro-IL-1β. This step ensures the inflammasome components are transcriptionally available but remains in an inactive state. Step 2 (Activation): Nigericin, a potassium ionophore, induces K+efflux, mitochondrial stress, ROS production, and the release of circulating cell-free mitochondrial DNA (ccf-mtDNA). These danger signals promote NLRP3 oligomerization, recruiting ASC and caspase-1, leading to IL-1β release. Additionally, BFE is hypothesized to modulate this pathway by interfering with mitochondrial stress, ROS production, or inflammasome assembly. Created with BioRender.com, Zachos,K (2025). *Maritan, Martina. (2019). Adapted from “NLRP3 Inflammasome Activation.” Retrieved from https://app.biorender.com/biorender-templates*

Following priming, nigericin, a microbial toxin produced by *Streptomyces hygroscopicus*, serves as a potent inflammasome activator by disrupting cellular ion homeostasis^42,43^. One of the key mechanisms by which nigericin activates NLRP3 is through potassium efflux (K+efflux)^44^. This toxin induces a rapid loss of intracellular K+, a critical upstream signal for NLRP3 activation. Reduced cytosolic K+concentrations facilitate NLRP3 oligomerization, an essential step in inflammasome assembly^44^. Additionally, nigericin disrupts mitochondrial homeostasis, leading to elevated ROS production and ccf-mtDNA release which serve as a secondary danger signal that amplifies NLRP3 activation^45^. Another potential mechanism of activation is lysosomal destabilization, where nigericin-induced cellular stress leads to cathepsin release into the cytosol, further promoting inflammasome assembly^42^.

Upon sensing these signals, NLRP3 undergoes oligomerization, facilitating the recruitment of ASC via CARD-CARD domain interactions^46^. ASC then binds pro-caspase-1, forming the NLRP3 inflammasome, a large multimeric complex essential for inflammatory cytokine maturation ^46^. Within the inflammasome, pro-caspase-1 undergoes cleavage into its active form, caspase-1, which in turn processes pro-IL-1β and pro-IL-18 into their biologically active forms. These cytokines are subsequently secreted into the extracellular space, where they contribute to amplified inflammation, immune cell recruitment, and lung tissue damage ^46^.

Our findings align with previous studies from our group and others showing that BFE and its main bioactive flavonoids (catechin, epicatechin, and epigallocatechin) can modulate NLRP3 activation through multiple mechanisms^21,24,32,47^. Previous studies have demonstrated that açaí extracts reduces ROS levels, preventing mitochondrial dysfunction, which is a key trigger for NLRP3 activation^48^. Our group has demonstrated via molecular docking studies that key flavonoids present at the biomatrix of BEF can bind to the NLRP3 pyrin domain, preventing its assembly^24^. Açaí extracts decreases the expression of IL-1β, caspase-1, and dsDNA release, reducing cell pyroptosis and inflammation^21,24,32,47^.These findings suggest that BFE directly modulates NLRP3 activation while also enhancing cellular homeostasis, making it a promising candidate for inflammatory lung disease management.

The activation of the NLRP3 inflammasome in lung epithelial cells and alveolar macrophages is highly relevant to acute lung injury (ALI) and acute respiratory distress syndrome (ARDS) ^49^. Additionally, in the context of lung transplantation, inflammatory responses that develop in the donor lungs is known to contribute to ischemia reperfusion injury, primary graft dysfunction and thereby worse patient outcomes post-transplant^50^. Elevated IL-1β and IL-18 levels contribute to excessive neutrophil infiltration, increased vascular permeability, and alveolar damage, ultimately exacerbating lung inflammation^51,52^. Furthermore, chronic NLRP3 activation has been implicated in the progression of COPD, asthma, and pulmonary fibrosis, conditions where persistent inflammation leads to airway remodeling and lung function decline^49,53^. Despite the clear role of NLRP3 in lung disease, there is currently no FDA-approved therapy directly targeting NLRP3. Existing treatments, such as corticosteroids and broad-spectrum anti-inflammatory drugs, provide symptom relief but fail to target the underlying inflammasome-driven pathology, also are associated with significant side effects. Synthetic NLRP3 inhibitors like MCC950 have shown promise but are limited by toxicity concerns, particularly hepatotoxicity, observed in clinical trials^9,10,28^. BFE presents a compelling alternative due to its: (1) efficacy in suppressing NLRP3 activation; (2) low toxicity and favorable safety profile and (3) cost-effectiveness as a natural bioactive extract. Given these advantages, BFE holds significant potential for clinical translation as a safe, accessible, and effective therapeutic option for managing chronic lung inflammation.

While our study provides *in vitro* evidence supporting BFE’s anti-inflammatory potential, several limitations must be acknowledged: (1) our findings are based on cell culture and lung organoid models. Future studies should include preclinical in vivo models to assess BFE’s systemic effects and pharmacokinetics; (2) while we demonstrated NLRP3 modulation, additional molecular studies (e.g., direct protein binding assays, transcriptomic profiling) are needed to fully elucidate BFE’s mechanism of action; (3) though BFE exhibited a favorable safety profile in vitro, long-term studies are required to confirm its safety in human applications.

In conclusion, this study highlights the therapeutic potential of BFE in suppressing NLRP3-driven lung inflammation. Our findings demonstrate that BFE effectively reduces ROS, pro-inflammatory cytokines, and inflammasome activation in both 2D lung cells and 3D lung organoids, with efficacy comparable to synthetic inhibitors like MCC950. Given the pressing need for safe and effective NLRP3 modulators, BFE emerges as a promising natural therapeutic candidate for managing chronic inflammatory lung diseases. Future research should focus on preclinical in vivo validation, dose optimization, and mechanistic studies to pave the way for potential clinical translation of BFE as a novel, cost-effective approach to inflammasome-targeted lung therapies.

## Acknowledgements

We would like to thank Alencar Machado’s group in Brazil for their expertise in formulating the BFE extract and Applied Organoid Core (ApOC) facility at the University of Toronto for their contributions and assistance in generating lung organoids. D.E.S.E.S. was supported by an EPIC Doctoral Award and a MITO2i Graduate Scholarship for this work.

