## Supplementary Figure 1 for "Bioactive Flavonoid Extract Suppresses NLRP3 Activation and Inflammation in 2D and 3D Lung Models"

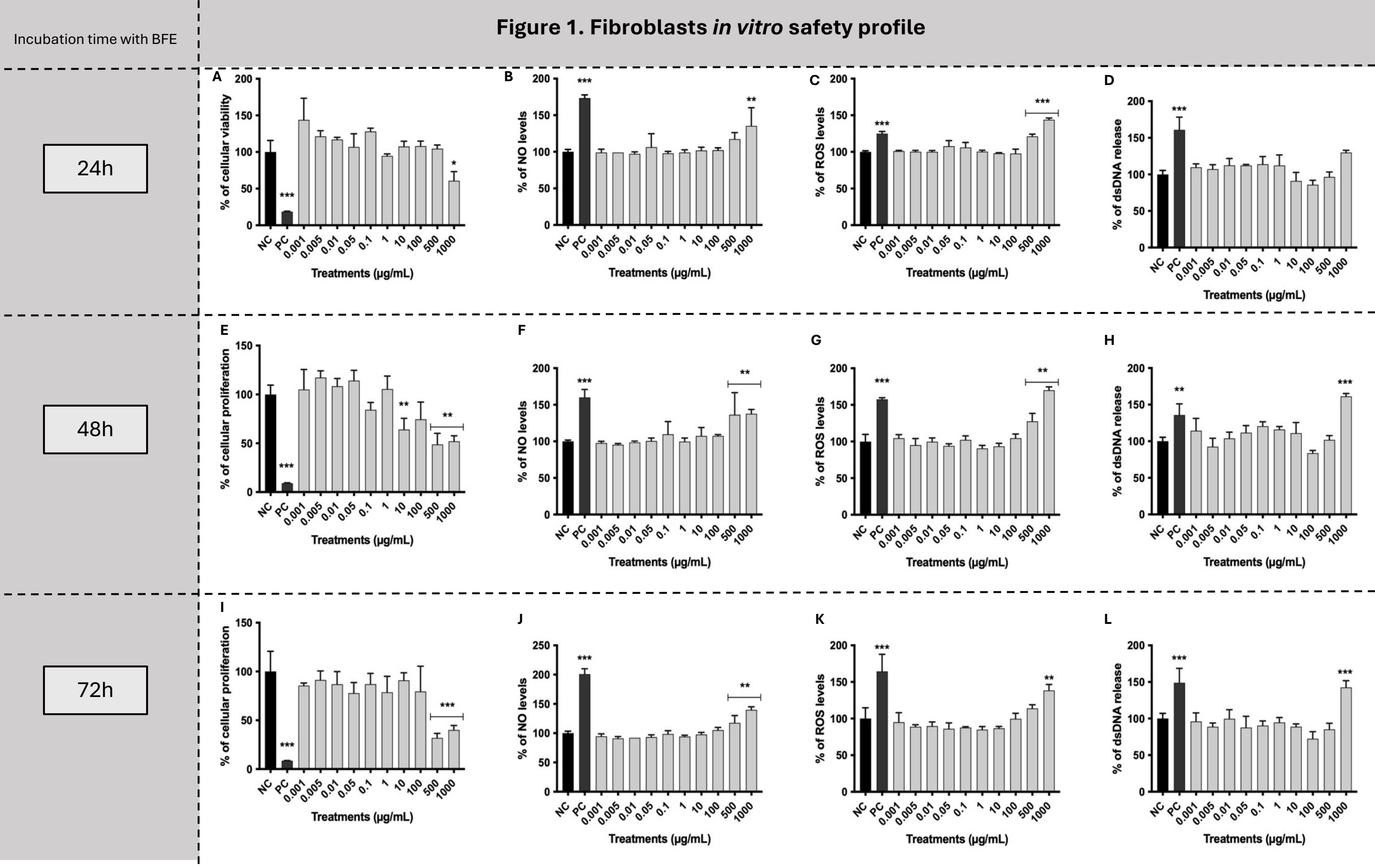


**Supplementary Figure 1.** Flavonoid enriched anti-inflammatory *agent* curve of concentrations - Toxicity evaluation in fibroblasts. Cells were treated with different concentrations of the BFE at 24, 48 and 72h of incubation. A, E and I) analysis of cellular viability (24h) and proliferation (48 and 72h) indexes by MTT assay; B, F and J) Indirect determination of NO levels; C, G and K) Qualitative measurement of ROS production by DCFH-DA assay; D, H and L) Release of dsDNA determination by using PicoGreen® fluorescent probe. Negative control (NC) are cells without any treatment; Positive control (PC) are cells exposed to 200 µM of $H_2O_2$ for MTT, DCFH-DA and PicoGreen assays and 10 µM of sodium nitroprusside for NO determination assay. Statistical analysis was performed by One-way Anova followed by Tukey *post hoc*. Results with p<0.05 were considered significant. *represents comparison to the negative control; *p<0.05; **p<0.01; ***p<0.001.
